# Water relations of *Calycanthus* flowers: hydraulic conductance, capacitance, and embolism resistance

**DOI:** 10.1101/182915

**Authors:** Adam B. Roddy, Kevin A. Simonin, Katherine A. McCulloh, Craig R. Brodersen, Todd E. Dawson

## Introduction

Flowers are developmentally and morphologically complex structures whose primary function is to promote sexual reproduction (Specht & Bartlett 2009). Coevolution with animal pollinators has long been considered the primary selective agent responsible for the many, diverse forms apparent among angiosperm flowers (Sprengel 1793, 1996; Fenster et al. 2004). Yet, floral adaptations to pollinators may not be as frequent as commonly considered, and non-pollinator agents of selection, such as the resource costs of building and maintaining flowers, may also influence floral form and function (Herrera 1996; Strauss & Whittall 2006). For example, the water costs of flowers can limit flower size and impede leaf function (Galen 1999; Galen et al. 1999; Lambrecht & Dawson 2007; Lambrecht 2013; Teixido & Valladares 2013, 2014). The requirement of maintaining water balance has led to coordinated shifts in hydraulic traits that vary systematically across the angiosperm phylogeny (Roddy et al. 2016) and with other pollination traits, such as floral lifespan (Zhang et al. 2017).

The amount of resources required to produce and maintain flowers and how and when these resources are supplied and provisioned to flowers may be highly variable among species and habitats (Bazzaz et al. 1987; Reekie & Bazzaz 1987a b; Teixido & Valladares 2014; Roddy et al. 2016). This variation could be due to a number of causes, including environmental conditions and differences in the functions performed by flowers (Galen 1993; Lambrecht & Dawson 2007; Roddy & Dawson 2012). For example, higher temperatures, increased evaporative demand, and nectar secretion can lead to higher carbon and water requirements of flowers (Patiño & Grace 2002; Patiño et al. 2002; De la Barrera & Nobel 2004). Furthermore, the mechanisms of water import may have shifted early in angiosperm evolution; early-divergent *Illicium* and *Magnolia* flowers are hydrated predominantly by the xylem (Feild et al. 2009a b) while some eudicot flowers are hypothesized to be hydrated by the phloem (Trolinder et al. 1993; Chapotin et al. 2003). Large variation in whole flower hydraulic conductance (*K_flower_*) and in the mechanisms of water import suggest that flowers may use a variety of mechanisms to remain turgid and functional during anthesis. Water may be continuously supplied by the xylem or by the phloem, or water imported early in development may be slowly depleted throughout anthesis. If water is continuously imported by the xylem, then embolism in flower xylem may detrimentally impact flower function (Zhang & Brodribb 2017). However, flowers tend to have high hydraulic capacitance (Chapotin et al. 2003), which can minimize water potential declines and help to physiologically isolate flower water status from changes in the water status of the rest of the plant. Flowers undoubtedly employ some combination of all these strategies, and there may be tradeoffs between these strategies, as there are in stems and in leaves (Brodribb et al. 2005; Meinzer et al. 2009; McCulloh et al. 2014).

In previous studies of *K_flower_* and other hydraulic traits (Roddy et al. 2013, 2016), flowers of the genus *Calycanthus* (Calycanthaceae; Zhou et al. 2006) consistently had traits associated with high rates of water supply (vein length per area; *VLA*), water loss (minimum epidermal conductance; *g_min_*), and high hydraulic conductance (*K_flower_*). These exceptional trait values, particularly for *C. occidentalis*, suggest that these flowers can transport substantial amounts of water, possibly even outpacing leaf transpiration. Using leaves as a comparison for understanding the water relations of these early-divergent flowers, we sought to characterize the diurnal patterns of gas exchange and water potential for flowers and, using pressure-volume relations, how this impacts flower drought tolerance. We focused our measurements on *C. occidentalis*, native to California, but include data for the other two species in the genus, *C. floridus*, native to the southeastern U.S., and *C. chinensis*, native to China.

## Materials & Methods

### Plant species, study site, and microclimate measurements

Between 5 and 25 May 2014, we studied three individuals of each of *C. occidentalis* and *C. chinensis* growing in a common garden at the University of California Botanical Garden. These data were supplemented with data collected in 2015-2017 from *C. floridus* growing in the U.C. Botanical Garden and the Marsh Botanic Garden, New Haven, CT. All statistical analyses were performed using R (v. 3.0.2; Team 2012).

Plants were kept well-watered throughout the study. The three *C. chinensis* individuals were growing in a more shaded microsite than the three *C. occidentalis* individuals. We characterized the microclimates and calculated vapor pressure deficit as (Buck 1981):

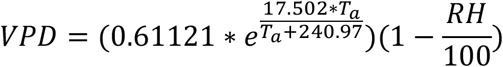

where *T_a_* and *RH* are air temperature (°C) and relative humidity (%), respectively. *Ta* and RH measurements were recorded every 5 minutes with a Hobo U23 (Onset Corp., Bourne, MA) data loggers that were housed in a covered, white, PVC T-shaped tube and hung 2 m off the ground within 100 m of both species.

At the U.C. Botanical Garden, *C. occidentalis* and *C. chinensis* flower beginning in May. During this time they have flowers in all stages of development. Flowers last for a few days, during which time those of *C. occidentalis* are entered by and provide shelter for venturous beetles (Grant 1950). Tepals tend to wilt and senesce starting at the tip moving towards the tepal base during anthesis. For all measurements, we sampled only newly opened flowers less than a day into anthesis.

### Diurnal measurements of gas exchange and water potential

We measured water vapor flux from entire flowers and parts of leaves of both species using an infrared gas analyzer equipped with a clear chamber (LI 6400 with LI 6400-05 conifer chamber, LI-COR Biosciences, Lincoln, NE). With this cuvette, leaf and flower temperatures were calculated based on energy balance, and the light level was not controlled. All measurements were made under ambient humidity and the reference CO_2_ concentration set to 400 ppm. At each time period on each plant, we measured at least one newly opened flower and a subtending leaf. We waited until fluxes had stabilized before recording 5 instantaneous measurements and subsequently averaging these. On 5 May 2014, we measured *C. occidentalis* at predawn (4:00 to 6:00 am) and every three hours after dawn (8:30 am, 11:30 am, 2:30 pm, 5:30 pm) and *C. chinensis* individuals at only predawn and midday (2:30 pm). Based on these data, the lowest daily water potentials and highest gas exchange rates occurred at midday (2:30 pm). Therefore, for subsequent measurements, we chose to sample only at predawn and midday and to sample multiple flowers per plant at these two time periods. In May 2015, predawn and midday water potentials of flowers, leaves, and stems were sampled again to supplement and corroborate measurements from 2014.

On the evening prior to gas exchange measurements, we covered one leaf subtending each flower in plastic wrap and aluminum foil so that this leaf could be used to estimate stem water potential on the subsequent day. Immediately after gas exchange measurements, the measured leaf and flower were wrapped in plastic wrap and excised with the adjacent foiled leaf and placed into a humidified plastic bag kept in a cool box and allowed to equilibrate for approximately 30 minutes. The balancing pressure was measured using a Scholander-style pressure bomb (SAPS, Soil Moisture Equipment Corp., Santa Barbara, CA, USA) with a resolution of 0.02 MPa. While inside the pressure chamber, leaves were kept covered by plastic wrap or plastic wrap and foil and flowers were wrapped in a plastic bag to prevent, as best as possible, desiccation inside the chamber during measurement. After water potential measurements, we transported the leaves and flowers to the lab and used a flatbed scanner and ImageJ (v. 1.47v) to estimate their surface areas, which were then used to recalculate gas exchange rates. Predawn gas exchange measurements for *C. occidentalis* leaves on 5 May are not included, however, because these leaves were misplaced before their areas could be measured.

From gas exchange and water potential measurements, we calculated flower and leaf hydraulic conductance based on Darcy’s law:

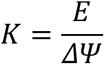

where *K* is the hydraulic conductance (mmol m^-2^ s^-1^ MPa^-1^), *E* is the transpiration rate (mmol m^-2^ s^-1^) and ΔΨ is the difference between stem and leaf or between stem and flower water potentials (ΔΨ_stem-leaf_ or ΔΨ_stem-flower_). This method assumes an approximate mass balance between the fluxes of liquid water into the structure (driven by ΔΨ) and water vapor loss from the structure (*E*).

### Pressure-volume analysis

We determined the relationship between Ψ and relative water content (RWC) of 4-5 whole flowers and leaves per species using repeated measures of mass and Ψ (Tyree & Hammel 1972). Flowering shoots were collected early in the morning on cloudy days. Mass was recorded immediately before and after each water potential measurement and subsequently averaged, and 10-12 measurements were made on each specimen as they slowly desiccated. Specimens were then oven-dried at 60^o^C for a week to determine dry mass. From the pressure-volume curves, we determined the Ψ and RWC at the turgor loss point (Ψ_TLP_ and RWC_TLP_) by a regression through at least five points of the linear part of the curve, the saturated water content per dry weight (SWC; g g^-1^) from the linear extrapolation to the x-intercept of the Ψ vs. water mass relationship normalized to dry mass, the modulus of elasticity from the slope of the relationship between turgor pressure and RWC above the turgor loss point, capacitance (MPa^-1^) and absolute capacitance (mol H_2_O kg^-1^ dry mass MPa^-1^) from the slope of the relationship between RWC and Ψ above the turgor loss point and the SWC. Extrapolation of these parameters was done in Microsoft Excel 2011, and differences between structures and species were analyzed using R. To test whether midday water potential declines are linked with hydraulic capacitance, we pooled data for species and structures and used standardized major axis regression to account for variance in both axes (the ‘sma’ function in the package *smatr*).

Stem hydraulic capacitance was calculated from water release curves of small chunks of small diameter (~1 cm) branches following previously published methods (McCulloh *et al.* 2014). Three shoots per species were sampled and five samples per species were used in the measurements. Samples were collected in the early morning, wrapped in wet paper towels, and kept refrigerated until analysis. All samples were vacuum infiltrated overnight in water. Excess water was removed from the samples by blotting them with paper towels, after which they were weighed and placed in screen cage thermocouple psychrometer chambers (83 series; JRD Merrill Specialty Equipment, Logan, UT, USA). Chambers were then triple-bagged and submerged in a cooler of water for 2-3 hours to allow equilibration between the sample and the chamber air. After equilibration, millivolt readings were recorded using a psychrometer reader (Psypro; Wescor, Logan, UT, USA). Samples were then removed from the chambers, weighed, and allowed to dry on the bench top for approximately 30 minutes before being returned to the chambers to repeat the measurements. The mV readings from the psychrometer reader were converted to MPa based on calibration curves from salt solutions of known water potentials (Brown & Bartos 1982). Samples were measured repeatedly until water potential values reached approximately -4 MPa, below which the psychrometers could not reliably resolve water potentials. Samples were oven-dried at 60^o^C for at least three days before weighing the dry mass. Relative water content (*RWC*) was calculated for each measurement and converted to relative water deficit (*RWD*) as 1 - *RWC*. The product of RWD and the mass of water per unit tissue volume at saturation (*M_w_*) yielded the cumulative mass of the water lost for each measurement interval. *M_w_* was calculated as:

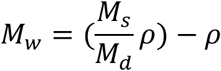

where ρ is wood density and *M_s_* and *M_d_* are the saturated and dry masses of the sample, respectively. The initial, linear phase of the plot of cumulative mass of water lost versus sapwood water potential gave the capacitance over the likely *in situ* physiological operating range of stem water potential (Meinzer et al. 2003, 2008). This regression of the initial linear phase was forced through the origin because of the physical impossibility of water being released at 0 MPa. How many of these initial points were used was similar to the method commonly used for analyzing pressure-volume curves of leaves. In this case, -1/Ψ was plotted against the amount of water released, and the number of initial points of this curve determined by adding points until the coefficient of variation declined. This final point was determined to be the inflection point on the moisture release curve, and capacitance was calculated as the slope of the regression between the origin and this inflection point on the plot of Ψ versus water released. Based on wood volume and density, hydraulic capacitance could be expressed in the same units as it is expressed for leaves and stems (mol H_2_O kg^-1^ dry mass MPa^-1^).

### Water loss rates and turnover times

At 11:45 am on 11 May, we excised three leaves and flowers per species to determine water loss rates. Cut surfaces were covered in petroleum jelly, and samples were weighed approximately every fifteen minutes on an electronic balance. Between measurements, samples were kept out of direct sunlight but not protected from ambient wind. Afterwards, specimens were scanned to determine surface area, and water loss rates were expressed as mmol m^-2^ s^-1^. Water loss rates measured in this way are a combination of stomatal and cuticular conductances. After excision and as tissue water potential declines, stomatal closure would reduce the relative contribution of stomatal conductance to overall water loss rates. To determine the effects of species and structure (flower vs. leaf) on water loss rates, we used a repeated measures ANOVA with species and structure within time as the error term.

Water turnover times were calculated from measurements of gas exchange and water potential as

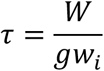

where *W* is the leaf or flower water concentration (mol m^-2^), *g* is the total conductance to water vapor (mol m^-2^ s^-1^), and *w_i_* is the mole fraction of water vapor inside the leaf or flower (mol mol^-1^). Because water content was not measured diurnally on the same samples measured for gas exchange, we estimated W from pressure-volume relations as the product of the relative water content, the saturated water content (g g^-1^), and the mass per area (g m^-2^). Measurements of mass per area for flowers were made solely on tepals, while pressure-volume parameters were calculated from entire flowers (tepals, pedicel, and receptacle), which makes our estimated of *W* for flowers lower than they may actually be. The relationship between *W* and Ψ derived from pressure-volume measurements (Figure S2) was used to estimate *W* based on diurnal measurements of Ψ. Two-phase linear relationships were used for the extrapolations, where different linear functions were fit to the relationship between *W* and Ψ above and below the mean turgor loss points.

### Stomatal responses to vapor pressure deficit

Steady-state stomatal responses of *C. occidentalis* flowers and leaves to VPD were determined using a Walz GFS-3000 gas exchange system. Flowering shoots were excised underwater in the morning and immediately recut underwater. Individual tepals or leaves were enclosed in the cuvette. Starting at approximately 0.7 kPa, VPD was increased in steps of approximately 0.3 kPa up to approximately 1.8 kPa for tepals and 2.8 kPa for leaves. Across this range of of VPD, gs declined to below 50% of its maximum for each structure. After each step change in VPD, *g_s_* was allowed to stabilize. Immediately after gas exchange measurements, the tissue enclosed in the cuvette was photographed and its area later measured using ImageJ to recalculate gas exchange rates. Data were compiled for replicates and non-linear curves fit and compared to linear curves using AIC.

### Characterizing embolism formation

In a separate experiment, we quantified the water potential at which embolism appeared in tepals using high resolution x-ray computed microtomography (microCT). In March 2015, flowering shoots of *C. floridus* were cut from plants growing at the U.C. Botanical Garden, recut underwater, and transported to the laboratory, where they were removed from water and allowed to slowly desiccate. Periodically, whole flowers were excised at the pedicel base and affixed in a custom, styrofoam holder that held 1-2 tepals in place. During microCT imaging, the entire flower was draped in a moist towel to minimize desiccation during the scan. The target tepals were imaged using continuous tomography at 24 keV while the sample was rotated from 0-180^o^. Images were captured by a camera (PCO EDGE; Cooke Corp., Romulus, MI, USA) with a 5x magnification Mitutoyo long working distance lens. Scans resulted in 1025 raw, two-dimensional projection images per sample, which were then reconstructed into tomographic slices using an Octopus Software ImageJ plugin (Institute for Nuclear Science, University of Ghent, Belgium). Immediately after microCT imaging of the samples, the scanned region of the tepals were excised and placed into thermocouple psychrometer chambers (83 series; JRD Merrill Specialty Equipment, Logan, UT, USA), which were then triple-bagged and placed into a 25^o^C water bath for approximately 4 hours, and the water potentials assessed as above. Scoring presence/absence of embolism in tepals was done visually by examining the tomographic slices for the presence of continuous air embolisms in veins. Logistic regression was used to calculate as a function of water potential the 50% probability of there being any embolism present, using the functions ‘glm’ and ‘dose.p’ in R (v. 3.0.2; Team 2012). In May 2015, the experiment was repeated for *C. chinensis* and *C. occidentalis*, but there were too few scans of high enough quality to accurately characterize embolism in these species, so we restrict this analysis to only *C. floridus*.

## Results

### Climate variation

The two main measurement days (5 May and 11 May) differed in their atmospheric conditions (Figure 1). On 5 May, temperature peaked at midday at 21^o^C, while on 11 May temperature peaked in the early afternoon at 25^o^C. These differences corresponded to different diurnal courses of VPD. On 5 May, VPD peaked at 1.09 kPa, while on 11 May, VPD peaked at 2.41 kPa.

**Figure 1.**
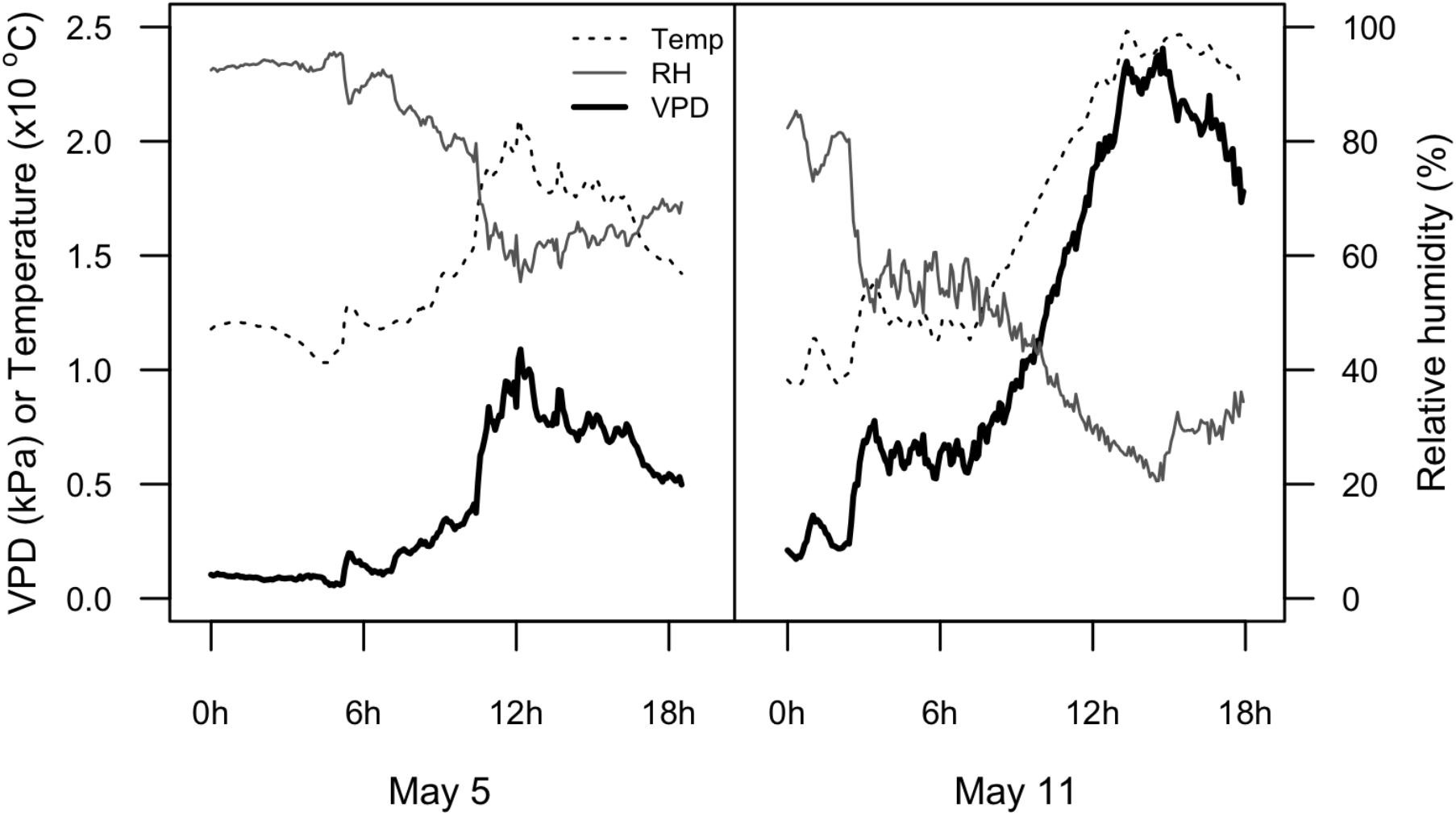
Diurnal variation in air temperature, relative humidity, and VPD during the two days of measurements.

### Pressure-volume relations

We used water release curves and pressure-volume relations to characterize the drought responses of stems, leaves, and flowers (Figure 2). For all species, water potentials at turgor loss were higher for flowers than for leaves (Figure 2a), with significant effects of species (F = 16.28, P < 0.001), structure (*F* = 55.74, *P* < 0.001), and the interaction of species and structure (*F* = 4.13, *P* = 0.032). The differences between structures, however, was significant only for *C. occidentalis* (*t* = 6.65, *P* < 0.001) and *C. chinensis* (*t* = 6.87, *P* < 0.01). Comparisons of *RWC_TLP_* were similar (Figure 2b), with a significant effect of species (*F* = 14.94, *P* < 0.001), structure (*F* = 22.98, *P* < 0.001), and the interaction of species and structure (*F* = 3.99, *P* = 0.034). The differences between structures were significant only for *C. occidentalis* (*t* = 3.69, *P* < 0.01) and *C. chinensis* (*t* = 7.77, *P* < 0.01). For all species, the saturated water content of flowers was higher than that of leaves (Figure 2c), with significant effects of species (*F* = 86.80, *P* < 0.001), structure (*F* = 1095.61, *P* < 0.001), and the interaction between species and structure (*F* = 33.76, *P* < 0.001). Difference between structures were highly significant for all species (*C. floridus*: *t* = 14.30, *C. occidentalis*: *t* = 17.62, *C. chinensis*: *t* = 26.51; all *P* < 0.001). Mass-specific hydraulic capacitance differed similarly (Figure 2d), with higher capacitance in flowers than leaves and significant effects of species (*F* = 22.89, *P* < 0.001), structure (*F* = 194.40, *P* < 0.001), and the interaction of species and structure (*F* = 15.05, *P* < 0.001). Differences between leaves and flowers were significantly different for all species (*C. floridus*: *t* = 7.20, *C. occidentalis*: *t* = 9.44, *C. chinensis*: *t* = 6.43; all *P* < 0.001). Based on the regression of water released versus stem water potential, stems of *C. floridus* had higher mass-specific hydraulic capacitance than either *C. chinensis* or *C. occidentalis* (102.8, 100.7, 88.0 mol kg^-1^ MPa^-1^, respectively; Figure 2d), and stems of all species had higher hydraulic capacitance than either leaves or flowers.

**Figure 2.**
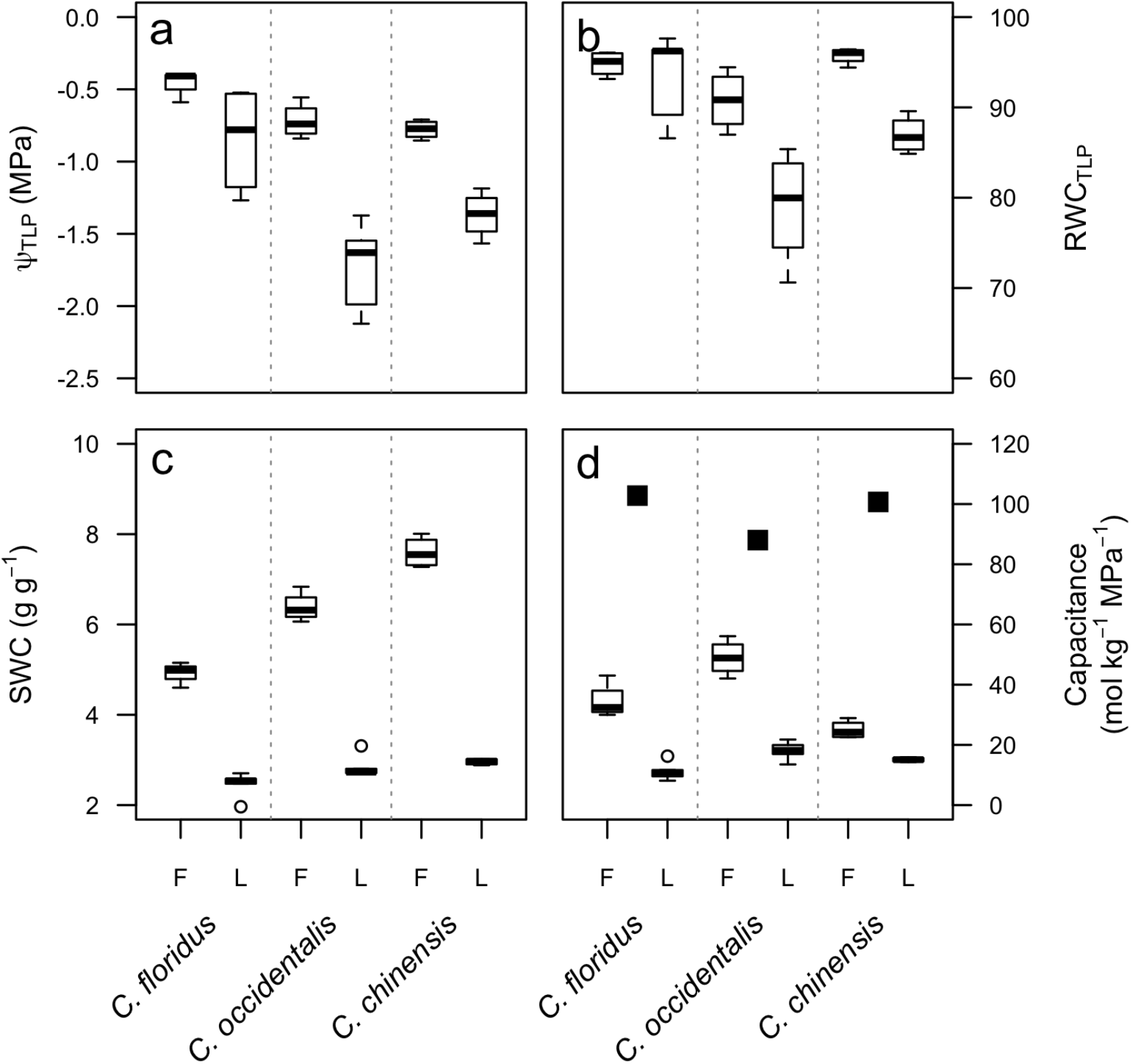
Parameters derived from the pressure-volume relationship for leaves (L) and flowers (F) of the three *Calycanthus* species: (a) water potential at the point of turgor loss, (b) relative water content at the point of turgor loss, (c) saturated water content relative to tissue dry mass, (d) hydraulic capacitance expressed as moles of water per kg dry mass per MPa. For comparison, stem capacitance values are plotted as filled squares in (d).

### Diurnal variation in water status and gas exchange

Diurnal variation in gas exchange and water potential highlighted differences between leaves and flowers. On 5 May 2014, measurements were made starting at predawn and continuing every three hours until 17:30 for *C. occidentalis* (Figure 3) and at predawn and midday for *C. chinensis*. Stomatal conductance and transpiration increased for both leaves and flowers throughout the day and peaked for both structures at 14:30. At this time, gs,leaf averaged 0.12 mmol m^-2^ s^-1^ and *g_s_*,flower averaged 0.09 mmol m^-2^ s^-1^. Similarly, transpiration peaked at 14:30, averaging 1.26 and 0.81 mmol m^-2^ s^-1^ for leaves and flowers, respectively. At all time points, gas exchange rates were higher for leaves than they were for flowers (Figure 3a,b). Similarly, leaf water potentials were always lower than both stem and flower water potentials (Figure 3c) with midday minimum water potentials of -0.93 and -0.77 MPa for leaves and flowers, respectively. The difference between stem and flower water potentials (ΔΨ *_stem-flower_*) varied little during the days, peaking at only 0.10 MPa at 14:30, in contrast to ΔΨ_*stem-leaf*_ which increased from 0.10 MPa at predawn to 0.25 MPa at midday. Measurements of water potential in May 2015 and 2016 showed similar patterns as measurements in 2014 for all three species (data for *C. floridus* in Figure S2). As a result of high *E* relative to the low ΔΨ, *K_flower_* was higher than *K_leaf_* at all time points except the last (Figure 3d). In the early afternoon (14:30), when *E* was highest, *K_flower_* averaged 14.45 mmol m^-2^ s^-1^ MPa^-1^ while *Kleaf* averaged 5.04 mmol m-2 s-1 MPa-1. These measurements of *Kf_lower_* based on transpiration rate were lower than the maximum *K_flower_* determined using the vacuum pump method except for the the peak measurements of *K_flower_* measured at midday (18.79 mmol m^-2^ s^-1^ MPa^-1^, solid, horizontal line and shading in Figure 3d; Roddy et al. 2016). In contrast, *K_leaf_* increased throughout the day, peaking in the early evening (17:30) at 10.24 mmol m^-2^ s^-1^ MPa^-1^ despite declines in both *g_s,leaf_* and *E_leaf_* after midday.

**Figure 3.**
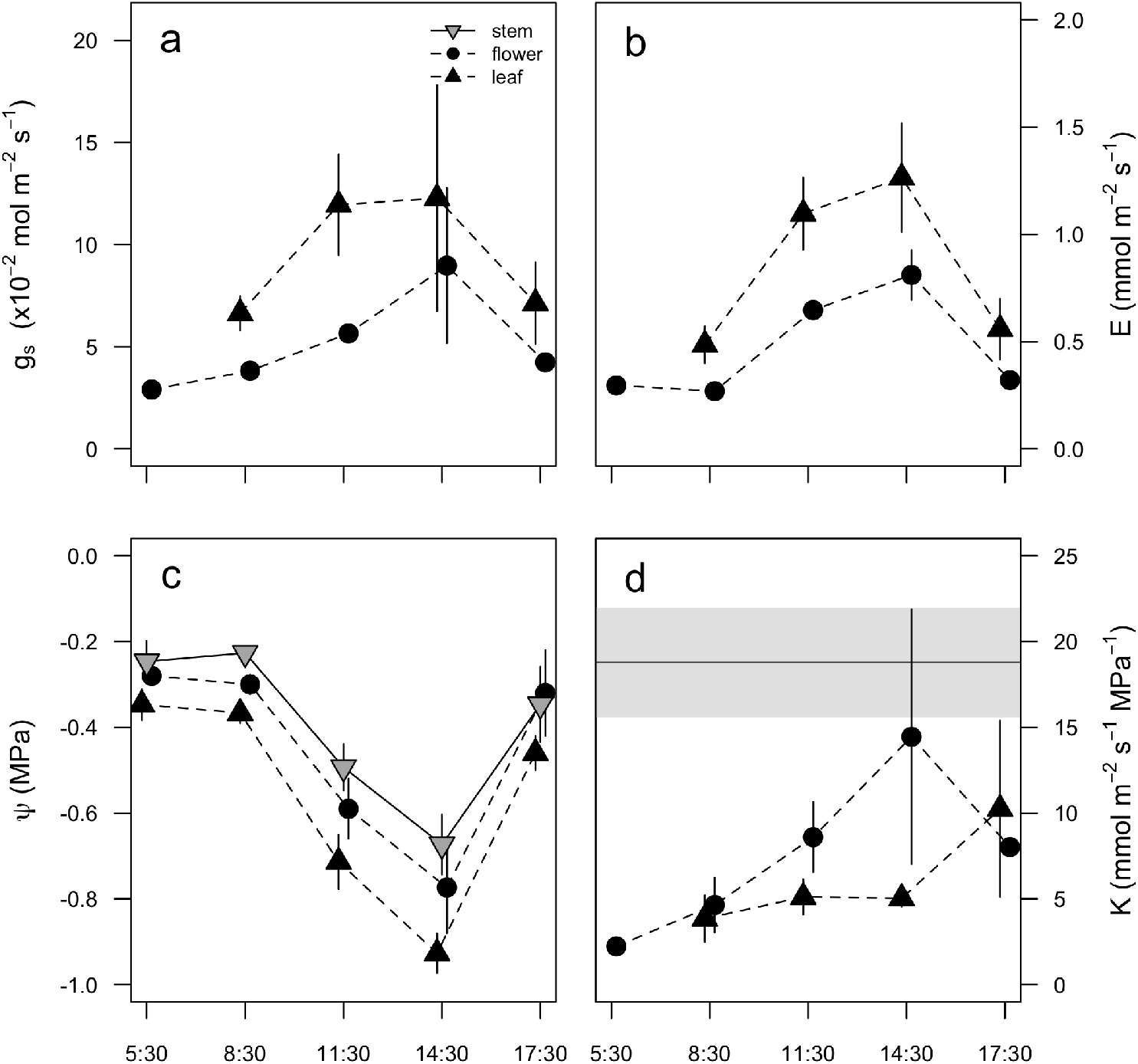
Diurnal measurements of (a) stomatal conductance, (b) transpiration, (c) water potential, and (d) hydraulic conductance on 5 May for *C. occidentalis*. Error bars represent standard error. Slight jitter in the horizontal axis as been added so that points and error bars do not overlap. In (d), the horizontal line and shading represent the average and standard error, respectively, of maximum *K*_*flower*_ for *C. occidentalis* reported in Roddy et al. (2016).

Because *g_s_* and *E* peaked for both leaves and flowers at 14:30, on 11 May 2014, we subsequently measured gas exchange and water potentials at only these times for both *C. occidentalis* and *C. chinensis* (Figure 4). At predawn and midday, *g_s,flower_* was higher than *g_s,leaf_* for both species. Similar to measurements on 5 May, Ψ_*flower*_ tracked changes in Ψstem throughout the day, such that ΔΨ *_stem-flower_* only increased slightly for *C. occidentalis*. In contrast, ΔΨ increased much more from predawn to midday for *C. occidentalis* leaves and for leaves and flowers of *C. chinensis* (Figure 4c). Consequently, *K_flower_* increased more than threefold for *C. occidentalis*, but slightly decreased for *C. chinensis*. On this day, too, *K_flower_* sometimes exceeded the average maximum value measured using the vacuum pump method for C. occidentalis flowers but not for C. chinensis flowers (Figure 4d). *Kleaf* increased throughout the day for *C. occidentalis* but even at midday was less than one-third the value of *K_flower_* for this species. *Kleaf* changed little throughout the day for *C. chinensis* (Figure 4d). Pressure-volume relationships had significant effects on these diurnal patterns of water potential, as maintaining a higher hydraulic capacitance significantly reduced midday ΔΨ, (R^2^ = 0.69, *P* = 0.04; Figure 8).

**Figure 4.**
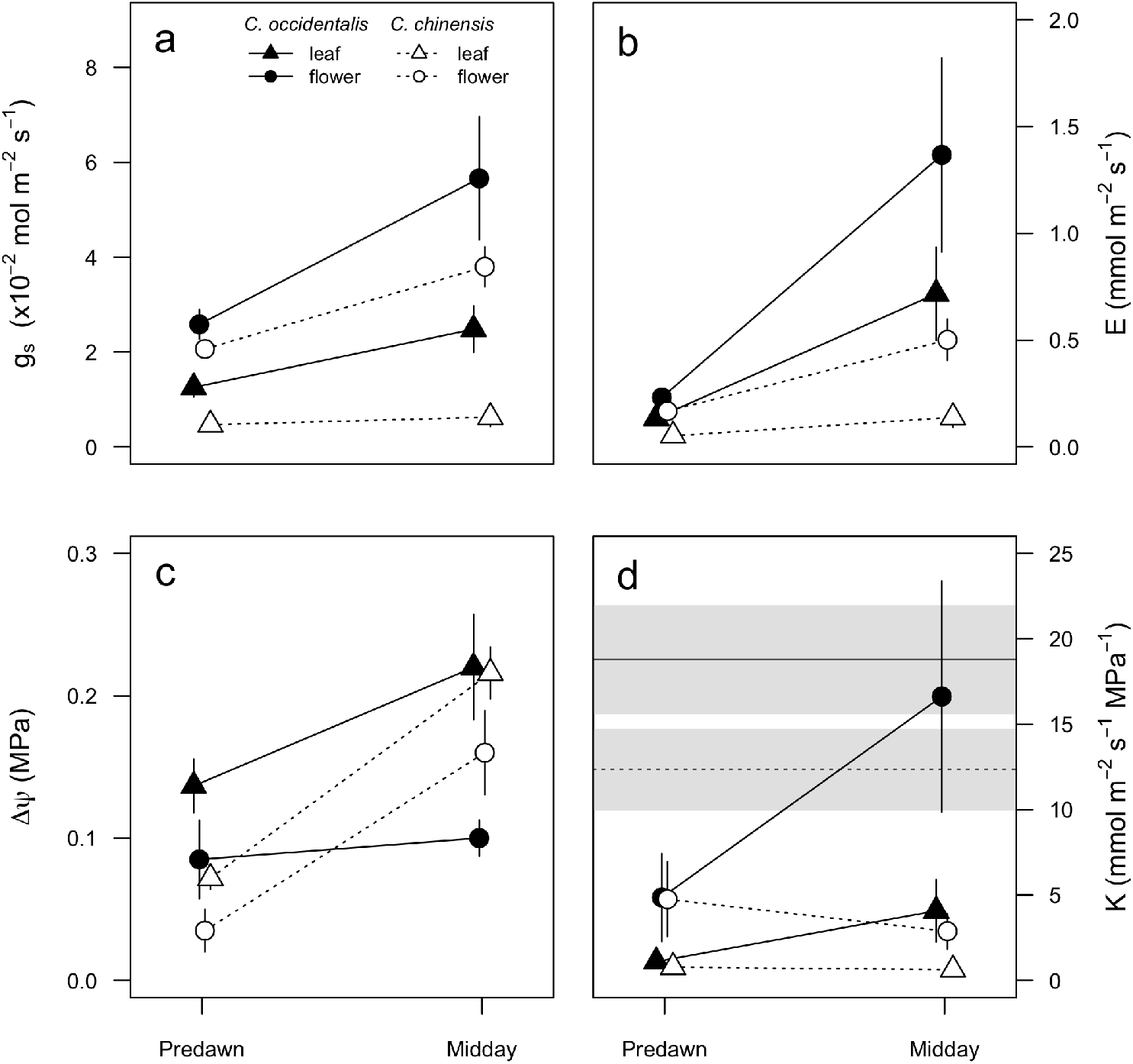
Predawn and midday measurements of (a) stomatal conductance, (b) transpiration, (c) water potential, and (d) hydraulic conductance of leaves and flowers for two *Calycanthus* species on 11 May. In (d), the horizontal lines and shading represent the averages and standard errors, respectively, of maximum *K*_*flower*_ for *C. occidentalis* (solid) and *C. chinensis* (dotted) from Chapter 3.

Average water loss rates differed significantly among species (*F* = 10.48, *P* < 0.01) and structures (*F* = 62.69, *P* < 0.001). Flowers lost water more rapidly than leaves, and both leaves and flowers of *C. occidentalis* (0.23 mmol m^-2^ s^-1^ and 1.14 mmol m^-2^ s^-1^, respectively) had higher water loss rates than those of *C. chinensis* (0.71 mmol m^-2^ s^-1^ and 0.10 mmol m^-2^ s^-1^). As a result, there was a significant interaction between species and structure (*F* = 7.87, *P* < 0.01; Figure 2). Flowers of both species had visibly wilted within 1 hour of excision.

### Response of stomatal conductance to vapor pressure

We characterized the gas exchange responses to VPD of leaves and flowers of *C. occidentalis* (Figure 5). There was no significant difference between maximum *g_s_* of leaves and tepals (120.8 ± 3.12 mmol m^-2^ s^-1^ and 152.8 ± 30.72 mmol m^-2^ s^-1^, respectively; *F* = 1.075, *P* = 0.35). However, the VPD at 50% of maximum gs differed among leaves and flowers. Tepal *g_s_* was 50% of maximum at 1.41 kPa, while leaf *g_s_* was 50% of maximum at 2.2 kPa.

**Figure 5.**
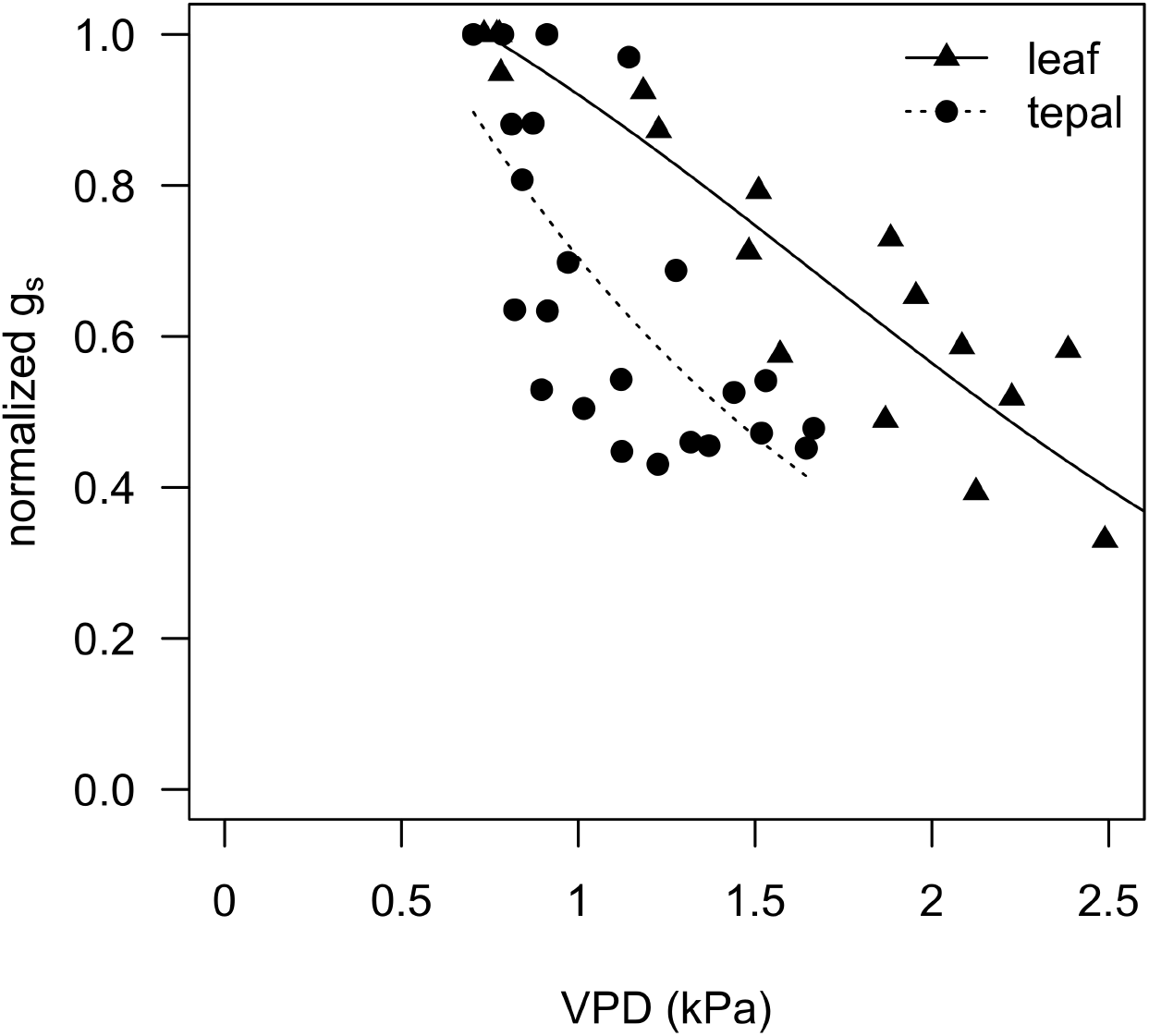
The responses of stomatal conductance for leaves and tepals of *C. occidentalis* to the vapor pressure difference driving transpiration. For each replicate, measurements of stomatal conductance were normalized to the maximum measured *g*_*s*_.

### Vulnerability to embolism in tepals

In the microCT experiment to characterize embolism, *C. floridus* ranged from -0.59 to -3.01 MPa, with no embolisms appearing until -2.03 MPa (Figure 6). The water potential at 50% probability of embolism was estimated to be -2.30 ± 0.22 MPa, significantly lower than the turgor loss point for *C. floridus* tepals (-0.45 ± 0.05 MPa). Interestingly, even at -3.01 MPa, only four conduits were embolized in the two tepals in this scan, suggesting that most conduits are even more resistant to embolism formation.

**Figure 6.**
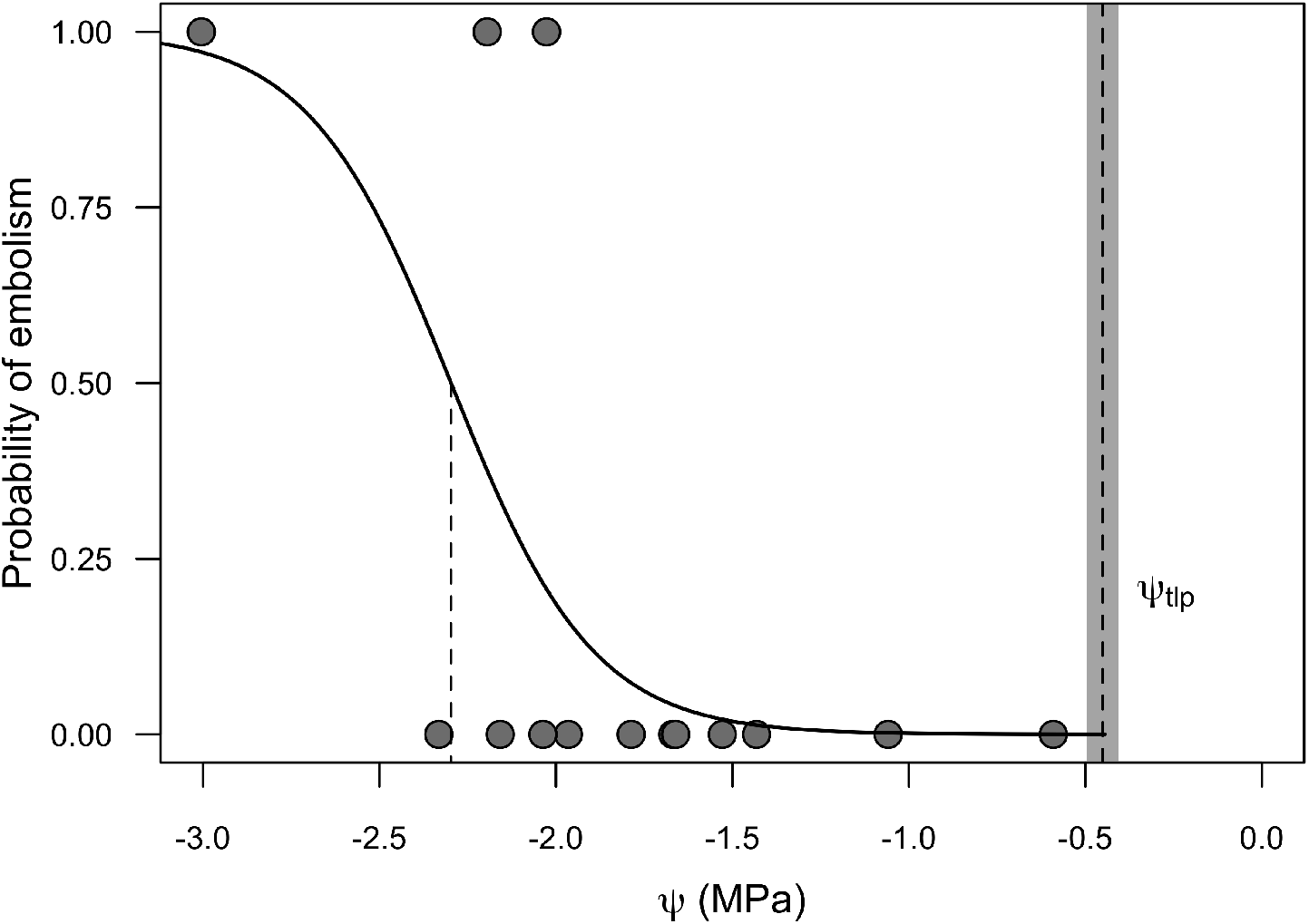
Vulnerability to embolism of *C. floridus* tepals as assessed by microCT imaging. The curve fit is indicated by a solid line with the water potential at 50% probability of embolism indicated by a dashed vertical line (-2.30 ± 0.22 MPa). The mean ± s.e. turgor loss point is indicated by the vertical dashed line and shading.

### Diurnal variation in water turnover times

Flower water turnover times (Figure 7) varied substantially throughout the day, from approximately 15 hours predawn to approximately 5 hours at midday during peak E for *C. occidentalis* on 5 May. Although leaf τ varied diurnally in a similar pattern, this variation was less dramatic. On 11 May, during which VPD was higher both predawn and throughout the day (Figure 1), predawn τ of flowers and leaves was higher than on 5 May (21 to 26 hours compared to 15 hours) but declined to similar values at midday, with little difference in midday τ between leaves and flowers of each species.

**Figure 7.**
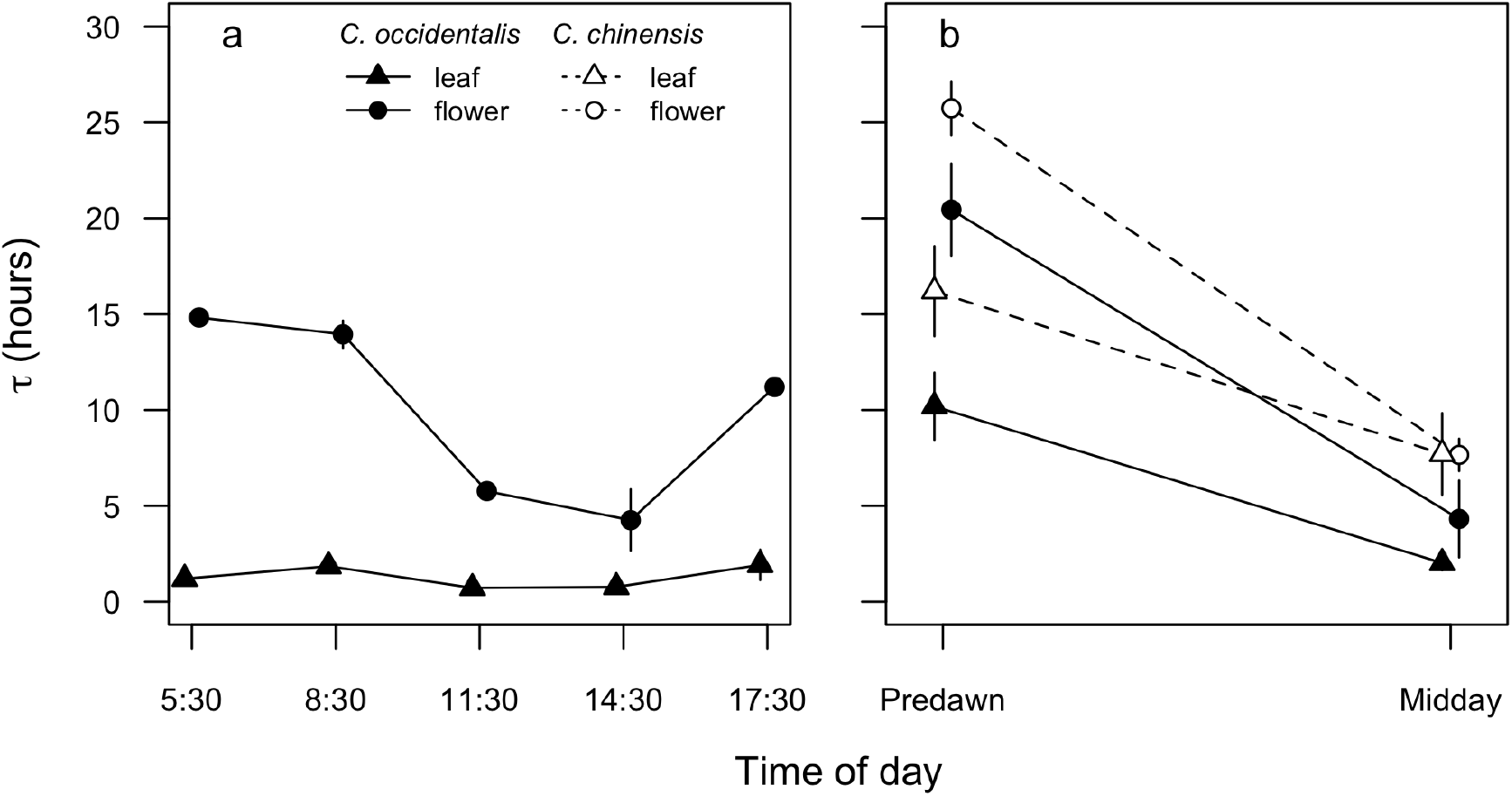
Diurnal variation in water turnover times (τ) calculated from diurnal measurements of gas exchange and water potential and from pressure-volume curve parameters (Figure S2). (a) Diurnal variation in τ for *C. occidentalis* on 5 May 2014. (b) Predawn and midday τ for *C. occidentalis* and *C. chinensis* on 11 May 2014. Points and vertical lines represent means and standard error.

**Figure 8.**
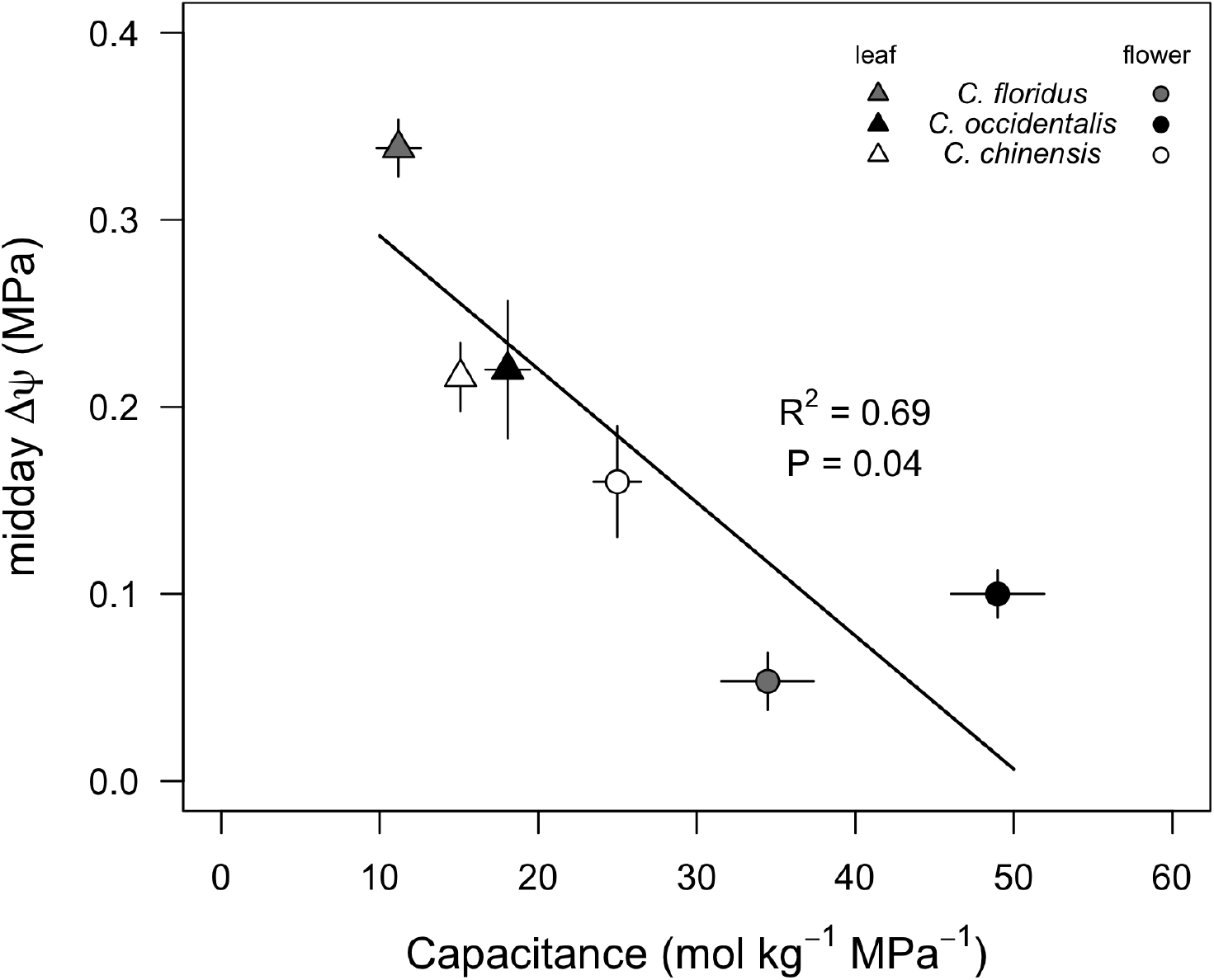
Tradeoff between hydraulic capacitance and midday water potential gradients (ΔΨ_stem-leaf_or ΔΨ_stem-flower_) pooled across species and days.

## Discussion

Many species flower under adverse conditions, such as when water is limiting. Maintaining flower function–turgid floral displays and receptivity–despite these resource demands is critical for sexual reproduction in many species. Even species native to mesic habitats, such as *Calycanthus* species, must support their flowers when atmospheric demand for water is high. Despite being well-watered, having high hydraulic capacitance, and having the highest *K_flower_* of any species measured, *C. occidentalis* flowers visibly wilted by midday when VPD was high.

### Regulation of transpiration

Gas exchange rates measured here for *Calycanthus* flowers were among the highest rates measured on flowers. Only tepals of avocado (*Persea americana*), another magnoliid, had transpiration rates comparable to those we measured for *Calycanthus* flowers (1.2-1.3 mmol m^-2^ s^-1^; Blanke & Lovatt 1993). Diurnal changes in *E* and *g_s_* observed here are similar to diurnal changes measured in other magnoliid flowers (Feild et al. 2009b) but notably different from estimates of *E* measured in eudicot *Cistus* species, in which *E* increased linearly with VPD (Teixido & Valladares 2014).

In some cases, transpiration from flowers can be higher than leaf transpiration. In both *Calycanthus* and *Persea*, *E_flower_* can be higher than *E_leaf_*, in contrast to measurements and estimates of *Eflower* in other species (Feild et al. 2009b; Roddy & Dawson 2012). Under cooler conditions, however, *Calycanthus* flowers also have lower *E* than their subtending leaves, and this variation is reflected in the functional responses to VPD and in the pressure-volume parameters. Although *Calycanthus* tepals had a higher maximum *g_s_* than leaves, tepal *g_s_* was more sensitive to VPD than leaf *g_s_* (Figure 5), probably because of a significantly higher Ψ_*TLP*_ (Figure 2a). Such high *g_s_* occured in *C. occidentalis* tepals despite their having lower stomatal densities than leaves (tepals: 14.31 mm^-2^, leaves: **XXXX**), suggesting that the high epidermal conductances of *Calycanthus* flowers, which are the highest among any flowers measured (Roddy et al. 2016), make significant contributions to total *E*. Indeed, *g_s_* reached its minimum and plateaued at 50% of maximum *g_s_* between approximately 1-1.5 kPa (Figure 5). This suggests that approximately half of maximum measured *g_s_* is due to epidermal conductance.

The high rates of *E* and *g_s_* measured in flowers may have important reproductive functions. Maintaining a high *g_s_*–and thus also *E* –would help to maintain floral temperatures below a critical, perhaps damaging, threshold temperature (Patiño & Grace 2002; Patiño et al. 2002). Because pollination in *Calycanthus* occurs when beetles crawl into and eat the tepals (Grant 1950), high *E* may provide a cool environment for beetles, protecting them from the hot, dry atmosphere. Higher *E* in flowers than leaves may also reflect prioritization of reproduction under conditions when even vegetative physiology suffers (Galen et al. 1999; Galen 2000). Given the relatively short lifespan of flowers and their relatively low surface area compared to leaves, at the whole plant level the water costs of flowers may still be small.

### Water balance and hydraulic autonomy of flowers

Maintaining water balance has likely been a critical constraint in floral evolution and diversification. The large, phylogenetically structured variation among species in *K_flower_* is coordinated with similarly structured variation in other anatomical and physiological traits associated with water supply and loss (Roddy et al. 2016). The high values of *K_flower_* for *Calycanthus* species–surpassing even *Kleaf* of many species–imply that they should be capable of transporting enough water during anthesis to meet their transpirational demands. Furthermore, *Calycanthus* flowers have significantly higher absolute hydraulic capacitance than leaves, which contributes to their lengthy water turnover times (Figures 2, 6). However, despite having both high *K_flower_* and high hydraulic capacitance, *Calycanthus* flowers were unable to prevent midday declines in psiflower that lead to turgor loss (Figures 2-5).

Maintaining high water potentials is important in order to avoid declines in water potential that would cause turgor loss and embolism formation and spread in the xylem. Our results indicate that *C. floridus* flowers have a large safety margin (~1.85 MPa) between turgor loss and embolism formation (Figure 6), and the rarity of embolized conduits even in the most desiccated tepals we measured (four conduits embolized at -3.01 MPa) suggests that *Calycanthus* tepals are at least as resistant to embolism as the species measured by Zhang and Brodribb (2017). Given that *Calycanthus* flowers lose turgor diurnally only on unusually hot days, the large safety margin between turgor loss and 50% probability of embolism suggests that embolism occurs only after flowers are no longer capable of rehydration. Autotrophic structures such as leaves must certainly avoid embolism to remain functional, but the effects of embolism formation on heterotrophic structures such as flowers is unclear. Maintaining a functional xylem pathway in flowers may matter little as long as it does not impede their ability to attract pollinators. Some flowers may still be able to attract pollinators even if petal water potential has declined past the point of turgor loss. Additionally, staggering flower opening over a period of days may be a bet-hedging strategy to prevent daily fluctuations in environmental conditions from yielding an entire season of flowers fruitless.

One way of maintaining high flower water potential would be to isolate them hydraulically from variation in stem water potential. Some have argued based on differences between leaf and flower water potential that flowers can be hydrated by the phloem rather than the xylem. Our results for *Calycanthus* flowers agrees with those from *Illicium anisatum* and *Magnolia grandiflora* that suggest that flowers are connected to the stem xylem during anthesis but disagrees with suggestions by others that flowers can remain more well-hydrated than stems (Trolinder et al. 1993; Chapotin et al. 2003; 2009a b). Data unequivocally showing the reverse Ψ gradients between stems and flowers thought to be indicative of phloem-hydration of flowers have been reported only for inner whorl tepals of *M. grandiflora* (Feild et al. 2009b). There is, however, more evidence available suggesting this for fruits. Reverse sap flow from developing mangoes into the stem occurs during the day as stem water potential declines (Higuchi & Sakuratani 2006). In tomatoes, most of the water was thought to be delivered by the phloem (Johnson et al. 1992), but recent evidence suggests otherwise (Windt et al. 2009). In grape pedicels, hydraulic resistance varies throughout development (Choat et al. 2009). This is driven by occlusion of the xylem, which is presumably important to allow the accumulation of sugars needed for grape berry ripening (Knipfer et al. 2015). Xylem discontinuity in the flower receptacle is another way to isolate floral structures from the stem xylem (Lersten & Wemple 1966). Both of these mechanisms could provide the spatial precision needed to differentially isolate heterotrophic and autotrophic floral structures (e.g. sepals vs. petals) and to generate the large water potential gradients observed between them (Trolinder et al. 1993). In *Calycanthus*, however, we have no evidence of xylem occlusion or discontinuity in the xylem pathway to the flower, particularly given the high values of *K_flower_* in the genus (Roddy et al. 2016). Like other magnoliid flowers, *Calycanthus* lacks a distinct calyx and corolla and has instead graded tepals. It may very well be likely that the evolution of a perianth differentiated into distinct autotrophic and heterotrophic structures also involved physiological differentiation in the mechanisms used to maintain water balance.

## Conclusions

The evolution of the flower was one of the hallmarks of angiosperm success. Despite the importance of flowers to reproduction, relatively little is known about their physiology and water relations. Here we show that *Calycanthus* flowers can have high stomatal conductance and transpiration, sometimes outpacing their subtending leaves, particularly during hot, dry conditions. Despite having among the highest hydraulic conductances of any flowers measured and despite reaching their maximum hydraulic conductance during peak atmospheric demand for water, *Calycanthus* flowers nonetheless can lose turgor and wilt. Despite stomatal closure and turgor loss, transpiration from flowers remains high due to high minimum epidermal conductance, suggesting that floral hydraulic architecture is not optimized for the extreme conditions encountered on particularly hot, dry days. Like other species of early-divergent angiosperm lineages with undifferentiated leaf-like tepals, *Calycanthus* flowers remain hydraulically connected to the stem xylem. Yet, these flowers rely heavily on hydraulic capacitance to minimize the water potential gradients with the stem because they cannot tolerate large water potential gradients. These complex dynamics between different hydraulic strategies could have important implications for our understanding of floral function, ecology, and evolution.

---

**Figure S1.**
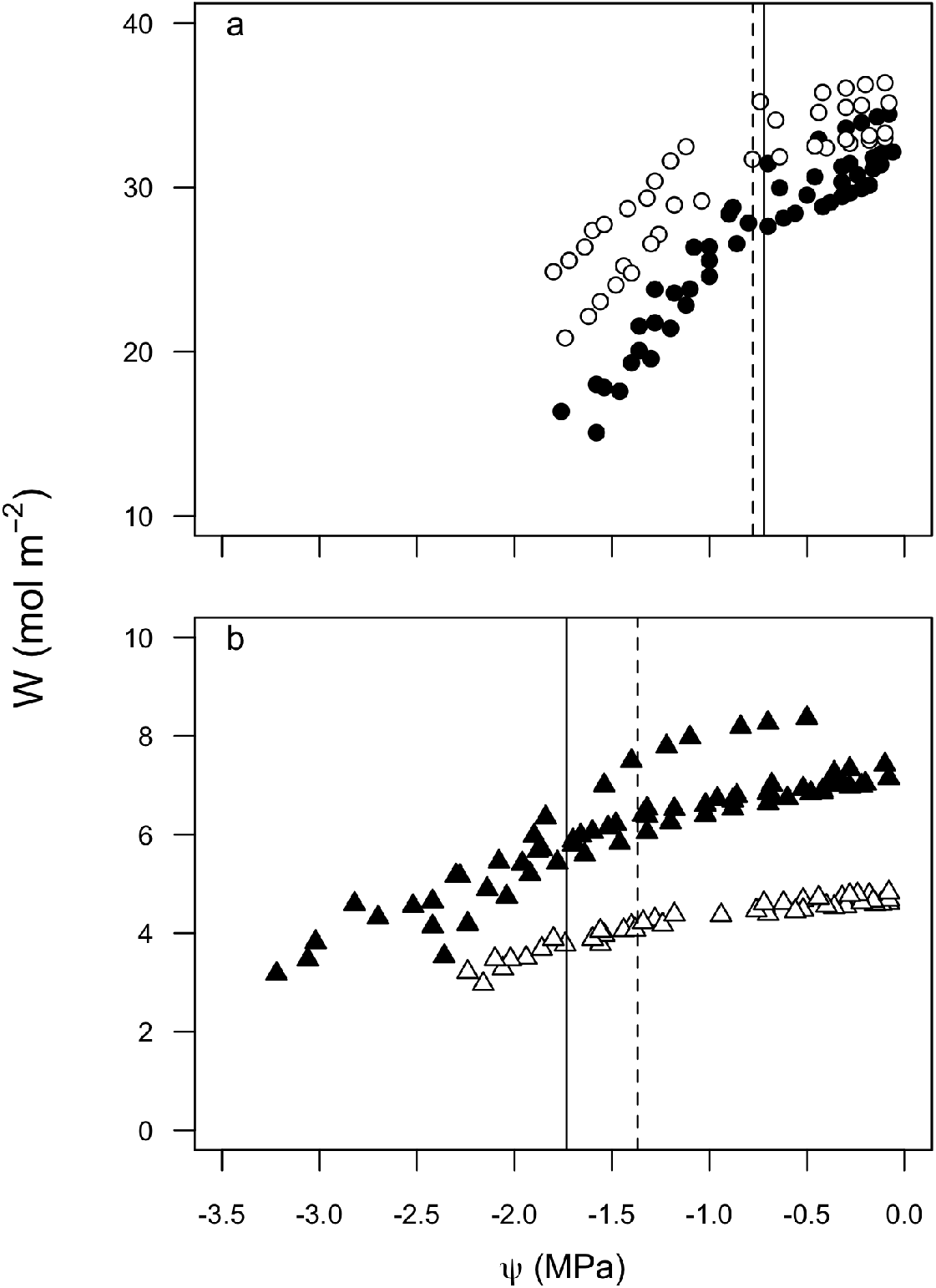
The relationship between molar water content (W) and water potential (Ψ) calculated from pressure-volume measurements. (a) Flowers of *C. occidentalis* (black) and *C. chinensis* (white) with their respective turgor loss points (solid and dashed lines, respectively). (b) Leaves of *C. occidentalis* (black) and *C. chinensis* (white) with their respective turgor loss points (solid and dashed lines, respectively). Note the different vertical axis ranges.

**Figure S2.**
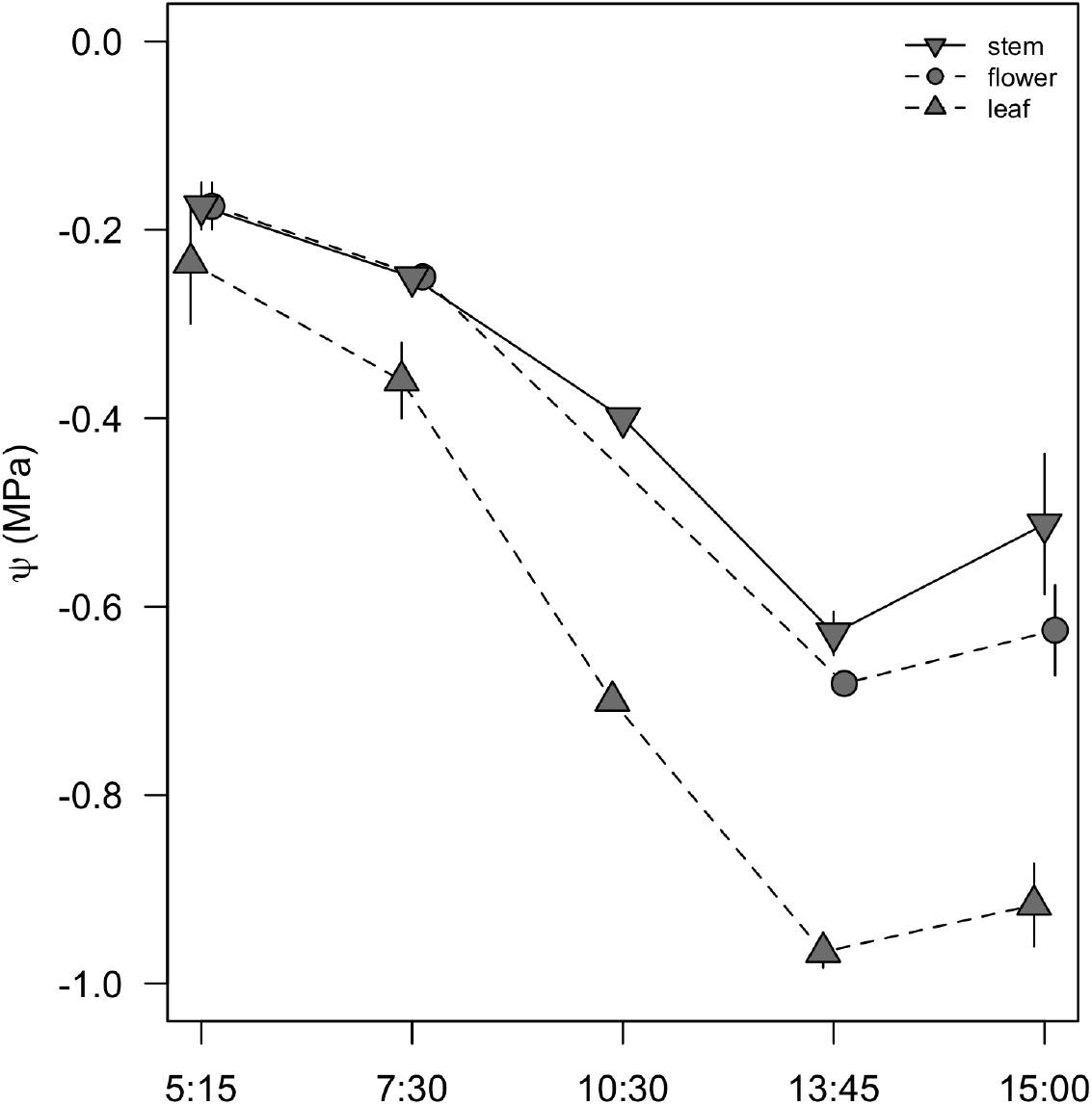
Diurnal water potentials for stems, leaves, and flowers of *Calycanthus floridus*.

